# Reciprocal regulation of vacuolar calcium transport and V-ATPase activity, and the effects of Phosphatidylinositol 3,5-bisphosphate

**DOI:** 10.1101/2020.05.22.111153

**Authors:** Gregory E. Miner, David A. Rivera-Kohr, Chi Zhang, Katherine D. Sullivan, Annie Guo, Rutilio A. Fratti

## Abstract

Yeast vacuoles are acidified by the V-ATPase, a protein complex comprised of the membrane embedded V_O_ complex and the soluble cytoplasmic V_1_ complex. The assembly of the V_1_-V_O_ holoenzyme is required for the transfer of H^+^ into the vacuole lumen for acidification. The assembly of the V_1_-V_O_ holoenzyme is stabilized by the lipid phosphatidylinositol 3,5-bisphospate (PI(3,5)P_2_) made by the PI3P 5-kinase Fab1/PIKfyve. The absence of PI(3,5)P_2_ leads to the dissociation of the V_1_ complex from the membrane. Separately, PI(3,5)P_2_ has been shown to modulate Ca^2+^ transport across the vacuole membrane during fission and fusion. Here we examined whether the regulation of H^+^ and Ca^2+^ by PI(3,5)P_2_ are interdependent. We show that modulating extraluminal Ca^2+^ concentrations inhibit V-ATPase activity. As extraluminal CaCl_2_ levels are raised, the activity of H^+^ pumping is reduced. Conversely, chelating free Ca^2+^ with EGTA stimulated vacuole acidification. Not only did Ca^2+^ levels affect H^+^ translocation, we also show that blocking V-ATPase activity inhibited Ca^2+^ transport into the vacuole lumen. Together, these data illustrate that Ca^2+^ transport and V-ATPase regulation are interconnected through the modulation of vacuolar lipid profiles.

**Summary Statement:** Here we show that Ca^2+^ and H^+^ transport across the vacuole membrane is reciprocally regulated and that it is linked to the production of Phosphatidylinositol 3,5-bisphoshpate.

## Introduction

The homeostasis of eukaryotic cells requires the active transport of elements across membranes against concentration gradients. In neurons, K^+^ accumulates in the cytoplasm and is released into the extracellular space leading to the depolarization of the membrane and import of Ca^2+^, which in turn triggers the fusion of synaptic vesicles with the plasma membrane to release neurotransmitters. Other gradients that are established include the accumulation of H^+^ in the lysosome to acidify the organelle and promote the activity of luminal hydrolases, and the storage of Ca^2+^ in the endoplasmic reticulum to regulate Ca^2+^ dependent signaling.

In *Saccharomyces cerevisiae*, H^+^ and Ca^2+^ ions are oppositely transported across the plasma membrane by the P2 type ATPase Pma1 and Cch1/Mid, respectively (Ariño et al., 2019, Locke et al., 2000). In yeast, both H^+^ and Ca^2+^ ions are mostly imported into the vacuole lumen. The V-ATPase pumps H^+^ into the vacuole lumen, while Ca^2+^ is transported into the vacuole by the Ca^2+^-ATPase Pmc1 and the Ca^2+^/H^+^ exchanger Vcx1 (Cunningham and Fink, 1994, Cunningham and Fink, 1996, Miseta et al., 1999). The vacuole contains Ca^2+^ at mM concentrations most of which bound to inorganic polyphosphate while a smaller pool is subject to further transport (Dunn et al., 1994). During osmotic shock, Ca^2+^ is released from the vacuole through the Transient Receptor Potential (TRP) channel ortholog Yvc1 (Dong et al., 2010). This activity requires the phosphatidylinositol 3-phosphate (PI3P) 5-kinase Fab1 and its production of PI(3,5)P_2_, which is also linked to vacuole fragmentation (Dong et al., 2010). Ca^2+^ efflux also occurs during vacuole fusion upon the formation of *trans*-SNARE complexes (Merz and Wickner, 2004). Unlike the fission pathway, vacuole fusion is inhibited by the production of PI(3,5)P_2_ after trans-SNARE complex formation that is linked to the inhibition of net Ca^2+^ efflux. (Miner et al., 2019a, Miner et al., 2020). PI(3,5)P_2_ lowers the observed net Ca^2+^ efflux through its activity on Pmc1. PI(3,5)P_2_ stimulates Ca^2+^ uptake by Pmc1. As short-lived PI(3,5)P_2_ is turned over, Pmc1 activity is reduced to basal levels of activity allowing SNARE-dependent Ca^2+^ efflux to be revealed. Thus, it appears that Fab1 activity could serve as a switch that promotes fission while inhibiting fusion through its effects on Ca^2+^ transport.

The regulatory role of Fab1 activity is not limited to Ca^2+^ transport and the fission/fusion switch. PI(3,5)P_2_ has been shown to regulate the V-ATPase and vacuole acidification through the direct physical interaction with the V_O_ subunit Vph1. This interaction stabilizes the assembly of V_1_-Vo complex to form the active V-ATPase (Banerjee et al., 2019, Li et al., 2014). In the Golgi, Vph1 is replaced by Stv1 and interacts with the compartment rich lipid PI4P instead of PI(3,5)P_2_, which is only made on late endosomes and lysosomes (Banerjee and Kane, 2017). In both instances, it is clear that specific phosphoinositides are essential for V-ATPase function.

The effects of PI(3,5)P_2_ on both V-ATPase function and Ca^2+^ transport suggests that these transport mechanisms could be interdependent. This notion is consistent with previous reports showing that inhibiting V-ATPase activity by deletion of components or treatment with Concanamycin A blocks the ability of Vcx1 to detoxify the cytoplasm after an increase in Ca^2+^ (Forster and Kane, 2000). Furthermore, Ca^2+^ transport is driven by a proton motive force, as ionophores such as CCCP and Nigericin block Ca^2+^ uptake (Ohsumi and Anraku, 1983).

In this study we examined the role of PI(3,5)P_2_ on the interdependence of Ca^2+^ and H^+^ transport during homotypic vacuole fusion. We demonstrate that Fab1 activity and its effects on Ca^2+^ transport inversely correlates with V-ATPase function. Separately we show that adding excess Ca^2+^ blocked V-ATPase activity, while chelating Ca^2+^ activated H^+^ pumping. Together, this study shows that H^+^ and Ca^2+^ transport is reciprocally regulated in a manner dependent on the lipid composition of the membrane.

## RESULTS

### Proton Influx is modulated by PI(3,5)P_2_

Recently, others showed that adding exogenous short chain dioctanoyl (C8) PI(3,5)P_2_ to purified yeast vacuoles augmented proton pumping (Banerjee et al., 2019). This was due in part to the ability of PI(3,5)P_2_ to stabilize the V_1_-Vo, an event was dependent on the interactions between the V_O_ subunit Vph1 and this lipid (Li et al., 2014). Our previous work showed that adding elevated concentrations of PI(3,5)P_2_ inhibited vacuole fusion at the hemifusion stage (Miner et al., 2019a). This was later linked to the ability of PI(3,5)P_2_ levels to affect Ca^2+^ transport across the vacuolar membrane (Miner et al., 2020). Based on these findings we hypothesized that PI(3,5)P_2_ may affect V-ATPase activity through modulating Ca^2+^ efflux from the vacuole lumen. To start, we performed acridine orange (AO) fluorescence quenching assays with isolated vacuoles from wild type yeast as well as strains expressing the kinase deficient *fab1*^EEE^ or the hyperactive *fab1*^T2250A^ mutations (Lang et al., 2017, Li et al., 2014). Because AO fluorescence decreases in acidic environments, it can serve as a measure of V-ATPase activity and pumping of H^+^ into the vacuole lumen. Vacuoles were incubated with AO for 600 seconds under conditions that promote fusion. This allowed the V-ATPase to operate leading to the quenching of AO. In **Figure 1A-B**, we show that the overproduction of PI(3,5)P_2_ by *fab1*^T2250A^ led to a pronounced increase in H^+^ pumping, as measured by increased AO quenching. This is in keeping with findings from the Kane lab and where they used of dioctanoyl (C8) PI(3,5)P_2_ in AO assays and found that H^+^ pumping was enhanced by this lipid but not by other C8 lipids. (Banerjee et al., 2019). We used C8-PI(3,5)P_2_ and found a similar effect (not shown). To show that the increased H^+^ transport activity by *fab1*^T2250A^ vacuoles was indeed due to the overproduction of PI(3,5)P_2_, we added the PIKfyve/Fab1 inhibitor Apilimod to reactions after 300 seconds of incubation (Cai et al., 2010, Dayam et al., 2015, Miner et al., 2020). Apilimod shifted the H^+^ pumping curve by *fab1*^T2250A^ vacuoles to wild type levels, indicating that the change in H^+^ pumping efficiency was due to the increase in PI(3,5)P_2_ **(Fig. 1B)**.

**Figure 1.**
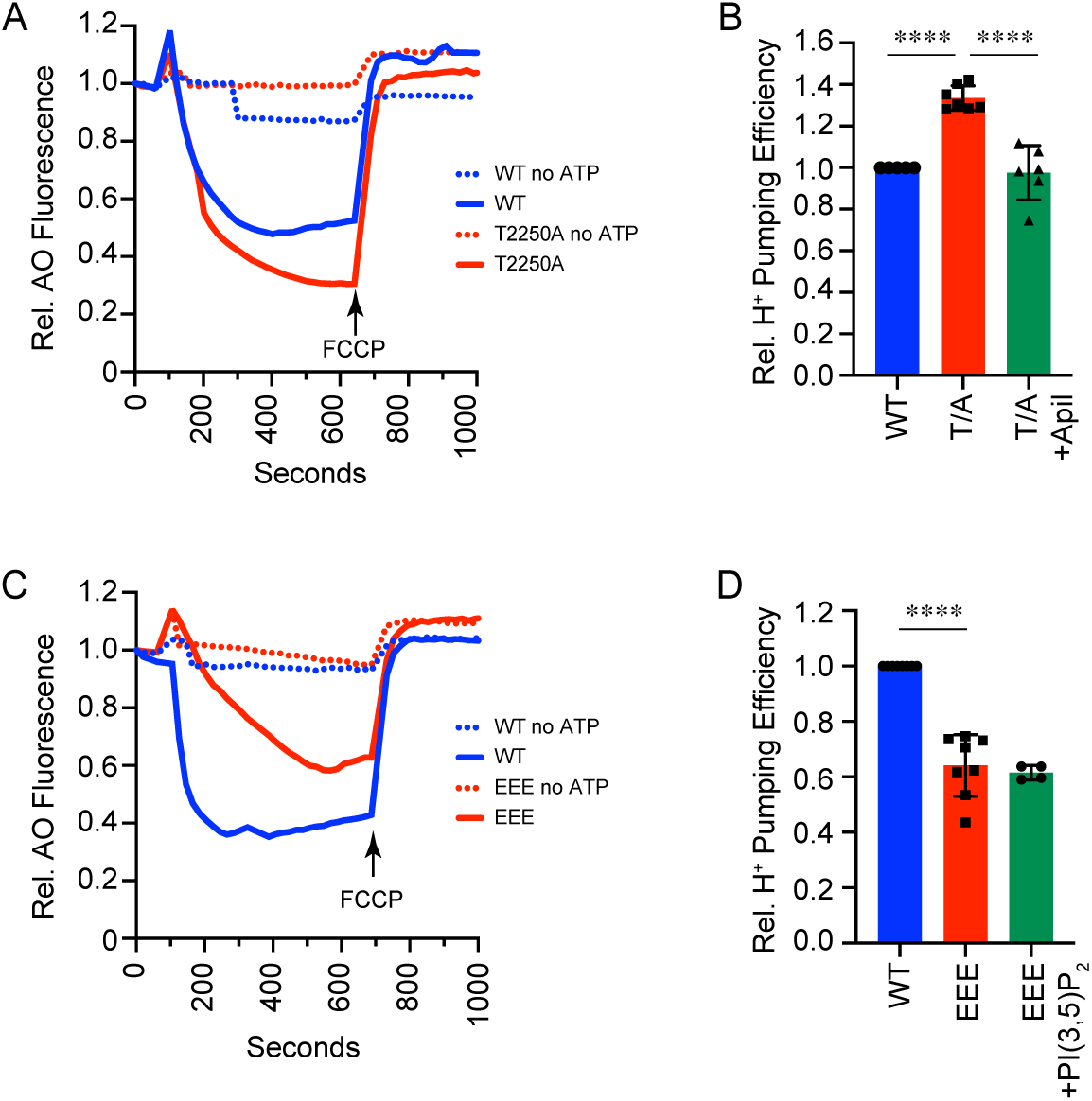
Regulatory lipids alter H^+^ translocation on isolated vacuoles. Vacuoles were used for proton pumping activity measured by acridine orange fluorescence quenching. Fusion reactions were incubated with or without ATP regenerating system added at 30 sec and the reactions were incubated for 300 sec. AO quenching was normalized to the initial fluorescence set to 1. **(A)** Wild type and *fab1*^*T2250A*^ vacuoles were compared in their efficiency to pump protons. In parallel *fab1*^*T2250A*^ vacuoles were incubated in the presence of 500 µM Apilimod. **(B)** Quantitation of multiple experiments in panel A. **(C)** Wild type and *fab1*^*EEE*^ vacuoles were compared in their efficiency to pump protons. Separately, *fab1*^*EEE*^ vacuoles were incubated 120 µM C8-PI(3,5)P_2_. **(D)** Quantitation of multiple experiments in panel C. After 700 sec reactions were supplemented with 30 µM FCCP was added to collapse the proton gradient and restore acridine orange fluorescence. Error bars are S.E.M. (n=3). \*\*\*\**p*<0.0001 (unpaired t-test).

While the increase in PI(3,5)P_2_ led to enhanced H^+^ pumping, we next asked if the lack of PI(3,5)P_2_ would reduce AO quenching. To this end we used vacuoles harboring the kinase deficient *fab1*^EEE^ mutant in the AO quenching assay. This showed that *fab1*^EEE^ vacuoles had attenuated H^+^ pumping **(Fig. 1C-D)**, which is in keeping with the destabilization of the V_1_-V_O_ complex when PI(3,5)P_2_ is absent (Lang et al., 2017). That said, enough V_1_-V_O_ is thought to remain on the vacuole to transport H^+^ into the vacuole lumen. To test if the difference was due to the absence of endogenous PI(3,5)P_2_, we supplemented the reaction with C8-PI(3,5)P_2_. Our data showed that supplementing *fab1*^EEE^ vacuoles with C8-PI(3,5)P_2_ failed in restoring wild type proton pumping **(Fig. 1D)**. Similarly, we previously found that the inhibition in hemifusion by *fab1*^EEE^ vacuoles was not rescued by C8-PI(3,5)P_2_ (Miner et al., 2019a). This does not mean that the lipid has no effect on *fab1*^EEE^ vacuoles. In a separate study we found that C8-PI(3,5)P_2_ restored wild type Ca^2+^ transport by *fab1*^EEE^ vacuoles (Miner et al., 2020). The lack of an effect was likely due to the altered sorting to the *fab1*^EEE^ vacuoles (Miner et al., 2019a). Together these data indicate that endogenous PI(3,5)P_2_ levels affect proton pumping. The lack of a complete inhibition of proton pumping by *fab1*^EEE^ vacuoles is in accord with data from the Botelho lab showing that *fab1*Δ vacuoles were able to acidify (Ho et al., 2015). They showed that *fab1*Δ vacuoles had a similar pH to wild type (pH 4.9) while vacuoles lacking the V_O_ subunit Vph1 have a pH of 6.1.

### Fab1 links V-ATPase activity to Ca^2+^ transport

In the experiments above we relied on either genetic mutations or exogenous sources to manipulate PI(3,5)P_2_ levels during AO quenching assays. In order to examine the rapid arrest of Fab1 function on V-ATPase activity we used Apilimod, which inhibits PI(3,5)P_2_ production by Fab1 during vacuole fusion (Miner et al., 2019a). In **Figure 2A-B**, we show that Apilimod completely inhibited AO quenching, suggesting that blocking PI(3,5)P_2_ production has an immediate effect on V-ATPase activity. The lack of an effect by DMSO treatment showed that the effect of Apilimod was not due to its solvent. Previously we found that Apilimod treatment enhanced Ca^2+^ efflux similar to the efflux profile seen with *fab1*^*EEE*^ vacuoles. We also found that treatment with the Ca^2+^ pump inhibitor Verapamil led to the accumulation of extraluminal Ca^2+^. Here we want to test whether the regulation of V-ATPase function by Fab1 occurs through modulating Ca^2+^ transport. To accomplish this, we treated vacuoles with Verapamil to block Ca^2+^ uptake by Pmc1 (Miner et al., 2020). We found that that Verapamil was as effective in blocking AO quenching compared to Apilimod, suggesting that an increase in extraluminal Ca^2+^ inhibited V-ATPase activity.

**Figure 2.**
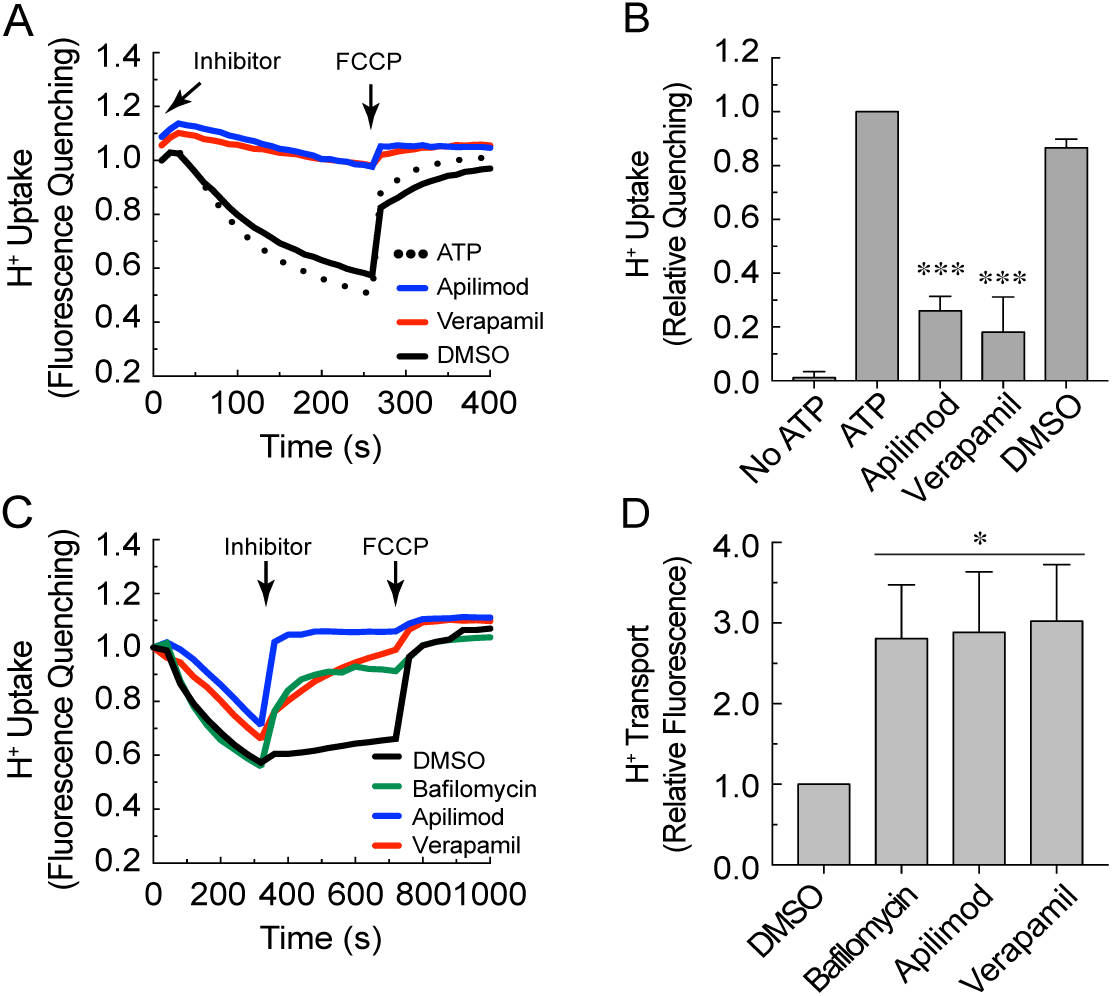
Apilimod blocks H^+^ transport. **(A)** Vacuoles were treated with Apilimod, Verapamil, DMSO in the presence of AO at the beginning of the reaction. AO fluorescence was measured for 250 seconds and FCCP was added at 260 seconds to collapse the H^+^ gradient (arrow). AO quenching was normalized to the initial fluorescence set to 1. **(B)** Average of multiple experiments represented in panel A. **(C)** Effect of late additions of Apilimod, Verapamil, Bafilomycin and DMSO on H^+^ transport (arrow, Inhibitor) added at 350 sec. FCCP was added at 720 seconds to collapse the H^+^ gradient (arrow, FCCP). **(D)** Average of H^+^ release in multiple experiments represented in panel C. Error bars are S.E.M. (n=3). \**p*<0.05, \*\*\**p*<0.001 (unpaired t-test).

To test whether inhibiting Fab1 at a later time affects the proton gradient, we added the reagent at 300 sec when AO quenching was complete. This showed a rapid release of H^+^ as shown by the de-quenching of AO. The rate at which it occurred was as quick as the effect of collapsing the gradient with FCCP. Further, the effects of Apilimod were similar to those seen with Bafilomycin A1 **(Fig. 2C-D)**. In addition, we found that adding Verapamil at this time point also lead to release of H^+^. Together these data suggest that Fab1 and PI(3,5)P_2_ production affect V-ATPase function through modulating Ca^2+^ transport. The effects of Apilimod and Verapamil were not due to leakage/lysis as these reagents do not inhibit vacuoles from fusing, which is inhibited by protonophores such as FCCP (Mayer et al., 1996).

To verify the effects seen with AO quenching, we used fluorescence microscopy and quinacrine staining with whole cells (Flannery et al., 2004, Miner et al., 2019b). Quinacrine accumulates and fluoresces in acidic compartments such as the yeast vacuole. As expected, untreated cells accumulated quinacrine in their vacuoles and fluoresced strongly **(Fig. 3, Buffer Control)**. When cells were incubated with CaCl_2_ there was a sharp decline in the number of quinacrine-labeled cells, indicating that an increase in cytosolic Ca^2+^ led to blocked V-ATPase activity. The increase in cytosolic Ca^2+^ was also accomplished with Verapamil to inhibit Pmc1 activity (Miner et al., 2020). Although Verapamil is known to inhibit voltage gated Ca^2+^ channels at low concentrations, it also inhibits Ca^2+^ ATPase pumps at higher concentrations. This includes mammalian SERCA (sarco/endoplasmic reticulum Ca^2+^ ATPase) and yeast Pmc1 (Miner et al., 2020, Paydar et al., 2005). As controls we used DMSO alone as well as the V-ATPase inhibitor Bafilomycin A1. DMSO treated cells stained with quinacrine, while those treated with Bafilomycin A1 showed little fluorescence. These data are in agreement with our data with isolated vacuoles and AO quenching.

**Figure 3.**
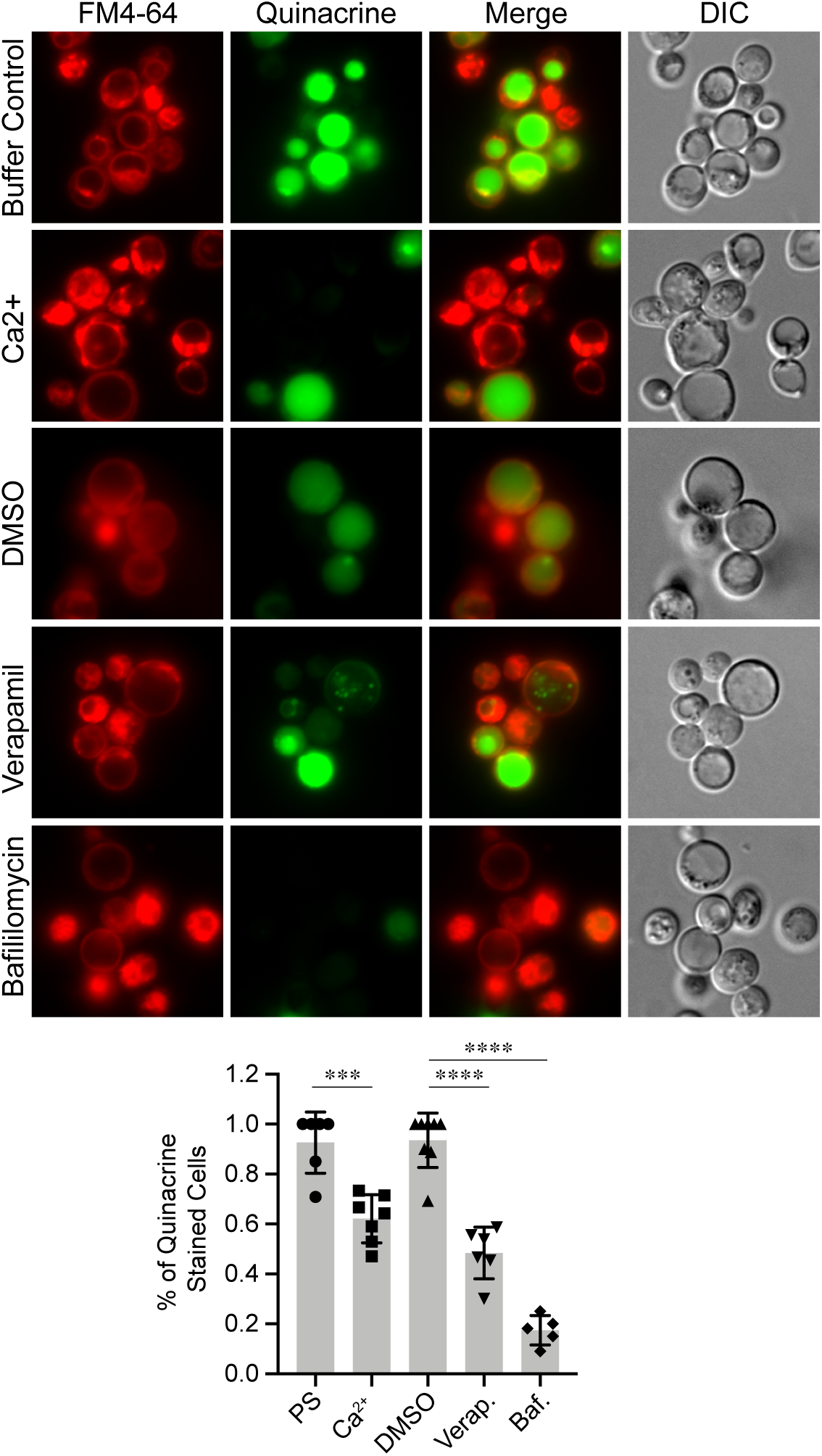
Quinacrine staining of yeast vacuoles. Log-phase BJ3505 cells were incubated with 200 µM quinacrine and 2 µM FM4-64 for 1h. DIC, differential interference contrast. Cells were additionally treated with DMSO, 100 nM Bafilomycin, 500 µM Verapamil or 250 µM CaCl_2_. Bar graph represents the average of cells strained with quinacrine. Error bars are S.E.M. (n=7 trials, 100 cells per condition, per trial. \*\*\**p*<0.001; \*\*\*\**p*<0.0001 (unpaired t-test).

### Ca^2+^ affects V-ATPase activity

Due to the link between the V-ATPase and Ca^2+^, we next tested if the direct addition or sequestration of Ca^2+^ would affect H^+^ pumping. First, we performed AO quenching assays with isolated vacuoles incubated with a dosage curve of CaCl_2_. This showed that Ca^2+^ at concentration of ≥100 µM abolished AO quenching **(Fig. 4A-B)**. In parallel we tested how vacuole fusion was affected by Ca^2+^. We found that AO quenching was partially inhibited at 100 µM CaCl_2_ with full inhibition at 250 µM. In comparison, vacuole fusion was not affected at these concentrations and only starts to be inhibited at concentrations of ≥500 µM CaCl_2_, which was in keeping with previous reports of vacuole fusion inhibition (Ungermann et al., 1999).

**Figure 4.**
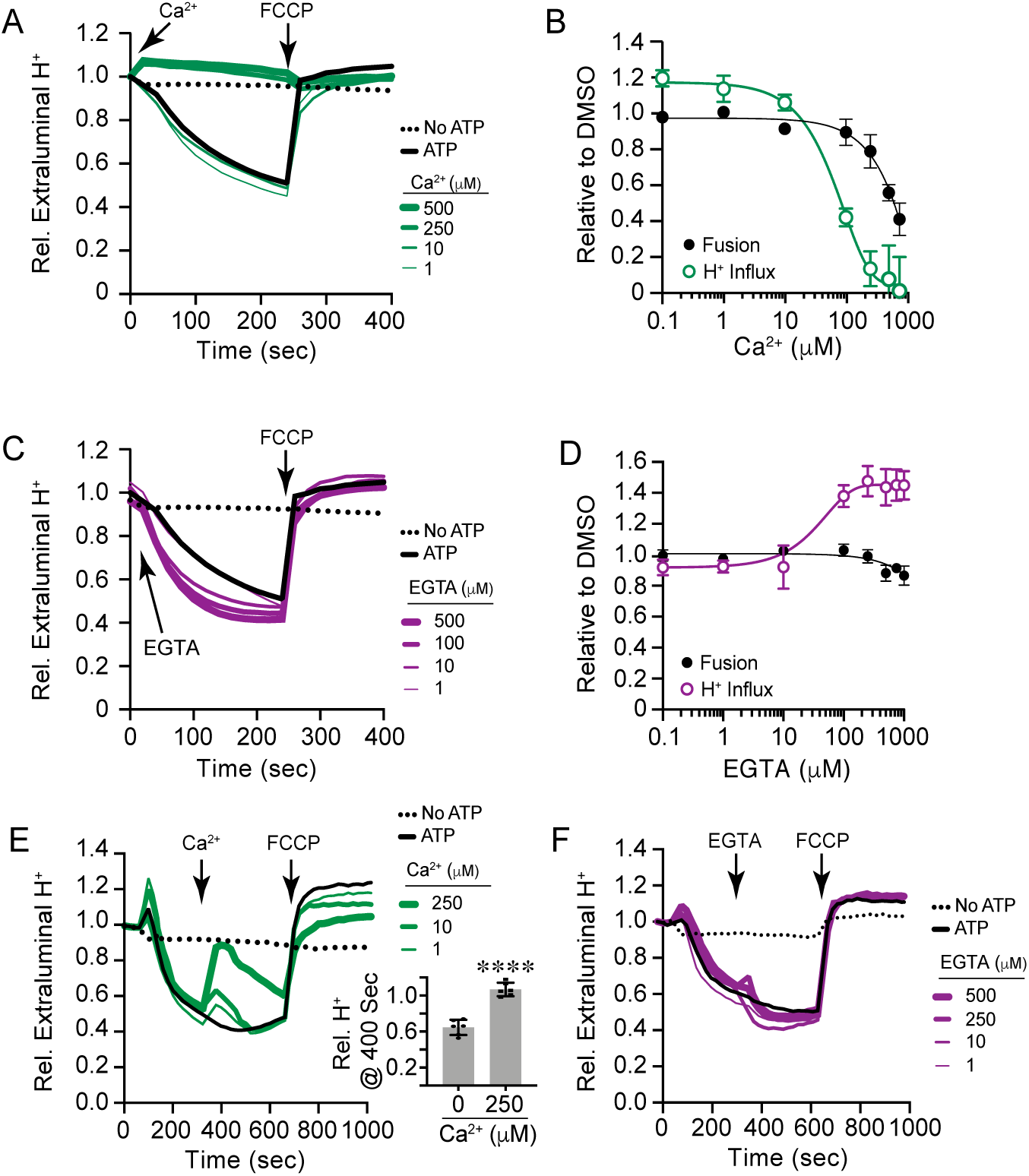
Ca^2+^ and H^+^ transport is reciprocally regulated during fusion. **(A)** H^+^ transport assays were performed in the presence of a dosage curve of CaCl_2_ at the indicated concentrations or PS buffer added at the start of the reactions. A separate reaction omitted ATP. Reactions were incubated for 240 sec after which FCCP was added (Arrow). AO quenching was normalized to the initial fluorescence set to 1. **(B)** Average of multiple experiments showing the effects of Ca^2+^ on AO quenching or vacuole fusion. **(C)** H^+^ uptake assay in the presence of a dosage curve of EGTA or PS buffer. **(D)** Average of multiple experiments showing the effect EGTA on H^+^ uptake and vacuole fusion. **(E)** Vacuoles were incubated for 300 sec at which point a curve of Ca^2+^ was added and further incubated for a total of 700 sec before addition of FCCP. Inset, average of extraluminal H^+^ of reactions treated with 250 µM Ca^2+^ or buffer alone. (F) AO quenching reactions were incubated for 300 sec after which a concentration curve EGTA was added and further incubate for a total of 700 sec before adding FCCP. Error bars are S.E.M. (n=3). \*\*\*\**p*<0.0001 (unpaired t-test).

**Figure 5.**
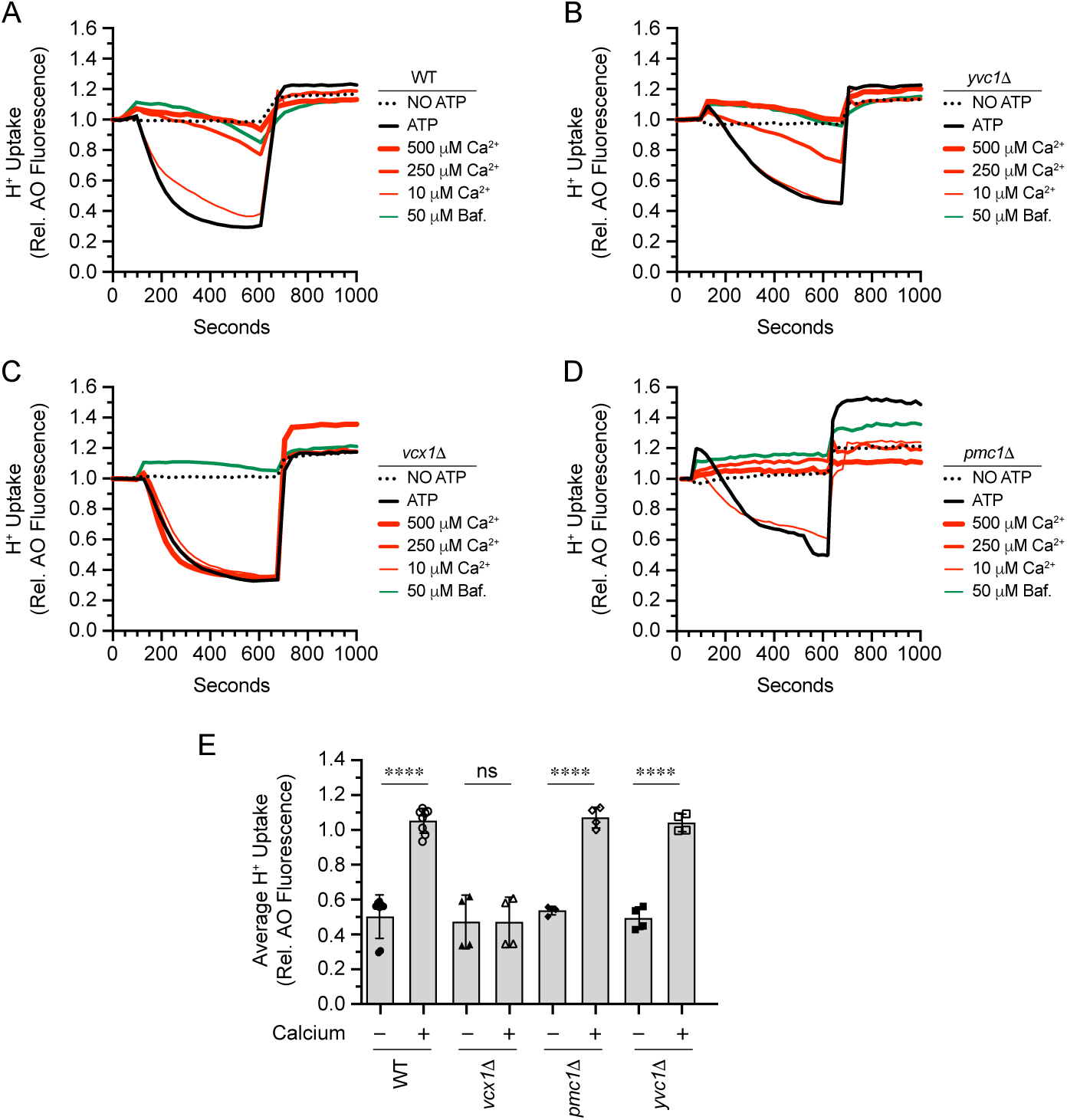
Effect of Ca^2+^ on V-ATPase activity with Ca^2+^ transporter mutants. Vacuoles from WT **(A)**, *yvc1*Δ **(B)**, *pmc1*Δ **(C)**, and *vcx1*Δ **(D)** were incubated with buffer, Bafilomycin A1 or a curve a CaCl_2_ for 600 sec. A separate reaction was performed in the absence of ATP. After ∼600 sec 30 µM FCCP to collapse the H^+^ gradient. AO quenching was normalized to the initial fluorescence set to 1. **(E)** Average of 4 experiments. Error bars are S.E.M. (n=3). \*\*\*\**p*<0.0001 (unpaired t-test); ns, not significant.

Based on the effect of adding CaCl_2_, we predicted that chelating Ca^2+^ would enhance V-ATPase activity. Thus, we performed AO quenching assays in the presence of a concentration curve of EGTA. As hypothesized, we found that EGTA enhanced AO quenching, suggesting that V-ATPase activity was increased **(Fig. 4C-D)**. While chelating Ca^2+^ enhanced V-ATPase activity, vacuole fusion was unaffected by EGTA. Together these data indicate that V-ATPase activity was affected by changes in Ca^2+^ levels. This was also in accord with our idea in which Fab1 activity correlates with increased V-ATPase function through modulating extraluminal Ca^2+^ levels.

We next asked whether the V-ATPase was sensitive to Ca^2+^ after H^+^ uptake had reached completion. AO quenching assays were started without exogenous Ca^2+^. After 300 sec, a concentration curve of CaCl_2_ was added to reactions and further incubated for a total of 700 sec before adding FCCP. These experiments showed that a rise in extraluminal Ca^2+^ resulted in AO dequenching in a dose-dependent manner suggesting that the V-ATPase was inhibited resulting in the accumulation of extraluminal H^+^ **(Fig. 4E)**. This was similar with findings by Cagnac et al., however in their study they added Bafilomycin A1 before adding Ca^2+^ preventing a direct comparison (Cagnac et al., 2010). The release of H^+^ was temporary as the V-ATPase regained activity once the excess Ca^2+^ was taken up. The release of H^+^ was likely due to the activity of Ca^2+^/H^+^ exchanger Vcx1. It also likely that the temporary inhibition of H^+^ uptake revealed H^+^ release from other sources including the Na^+^(K^+^)/H^+^ exchanger Nhx1 (Brett et al., 2005, Qiu and Fratti, 2010). It should be noted that AO quenching was restored when 250 µM Ca^2+^ was added late in the reaction **(Fig. 4E)**, while addition at the beginning prevented AO quenching for the duration of the experiment **(Fig. 4A)**. This suggests that once vacuoles have taken up H^+^ to equilibrate the system the vacuoles can recover from the Ca^2+^ spike over time to reactivate V-ATPase function. Alternatively, it is possible that Vcx1 is more active after the initial uptake of H^+^ to remove the excess Ca^2+^ and restore V-ATPase function.

Based on the effects of adding Ca^2+^ late in the reaction, we asked if adding EGTA would further enhance H^+^ uptake when added late. Unlike the effects of excess Ca^2+^, the addition of EGTA at 300 sec did not lead to further H^+^ uptake. While there was a consistent increase in AO quenching, it was not significant **(Fig. 4F)**. We attribute the absence of a pronounced effect to lack of extraluminal Ca^2+^ after 300 sec of incubation. Ca^2+^ uptake is typically completed between 300 and 500 sec (Miner et al., 2016, Miner et al., 2017, Miner et al., 2019a, Miner et al., 2020).

### Ca^2+^ inhibits V-ATPase activity through Vcx1

To determine which Ca^2+^ transporter was linked to the effects on V-ATPase activity we used a panel of deletion strains lacking the Ca^2+^ exporter channel Yvc1, the Ca^2+^-ATPase pump Pmc1, and the Ca^2+^/H^+^ exchanger Vcx1. The yeast vacuole can only take up Ca^2+^ through Pmc1 or Vcx1, therefore the deletion of one can function as a reporter for the other. Wild type and transporter mutant vacuoles were used in AO quenching assays in the presence of a CaCl_2_ concentration curve. Vacuoles lacking Yvc1 and Pmc1 were equally sensitive to excess Ca^2+^ compared to the wild type. The results with *yvc1*Δ vacuoles were as predicted since the TRP channel is an exporter of luminal Ca^2+^. The sensitivity seen with *pmc1*Δ vacuoles suggests that Vcx1 mediated transport of Ca^2+^ is linked to the effect on V-ATPase function. When *vcx1*Δ vacuoles were tested it showed that they were resistant to the excess Ca^2+^ as AO quenching was unaffected. This suggests that Pmc1 function was not associated the block in V-ATPase function. This was unexpected as we have shown that Pmc1 is important in Ca^2+^ transport in response to changes in PI(3,5)P_2_ levels and interacts with the V_O_ component Vph1 (Forster and Kane, 2000, Miner et al., 2020). However, these findings are in accord with those of Förster & Kane showing that Vcx1 was chiefly responsible for responding to Ca^2+^ spikes (Forster and Kane, 2000).

### Effects of altering V-ATPase function on Ca2+ transport

Because Ca^2+^ affects V-ATPase activity, we next asked if the reverse was also true. To achieve this, we used chemical inhibitors of the V-ATPase and monitored Ca^2+^ transport. In **Figure 6A-B** we show the effects of using a concentration curve of Bafilomycin A1 on Ca^2+^ transport. The transport of Ca^2+^ in and out of the vacuole was detected by the Ca^2+^ binding fluorophore Cal520 dextran conjugate, which fluoresces when bound to Ca^2+^ (Miner et al., 2020, Miner and Fratti, 2019). Adding Bafilomycin A1 at the beginning of the Ca^2+^ flux assay blocked Ca^2+^ uptake in a dose dependent manner. At ≥100 nM, Bafilomycin completely abolished Ca^2+^ uptake, indicating that V-ATPase function affects the Ca^2+^ transport machinery. As controls we used anti-Sec17 IgG to prevent trans-SNARE pairing dependent Ca^2+^ efflux (Merz and Wickner, 2004). In a separate reaction we omitted ATP to prevent Ca^2+^ uptake. Finally, we used DMSO to examine if the solvent for Bafilomycin had a significant effect on Ca^2+^ transport. Separately, we tested Bafilomycin A1 against vacuole homotypic fusion and found that curve for fusion inhibition followed the same trend as the one for Ca^2+^ uptake, albeit the effect on fusion was moderate and only reduced it by ∼40% **(Fig. 6B)**. To verify that the effect of Bafilomycin A1 was specific for the inhibition of V-ATPase activity, we used Concanamycin A. Adding Concanamycin A to vacuole reactions had a similar effect as Bafilomycin in that Ca^2+^ uptake was fully inhibited **(Fig. 6C-D)**. Concanamycin A also reduced fusion by 50% at 100 nM **(Fig. 6D)**. Together these data suggest that the Ca^2+^ transport machinery and the V-ATPase are interdependent and reciprocally regulated one another.

**Figure 6.**
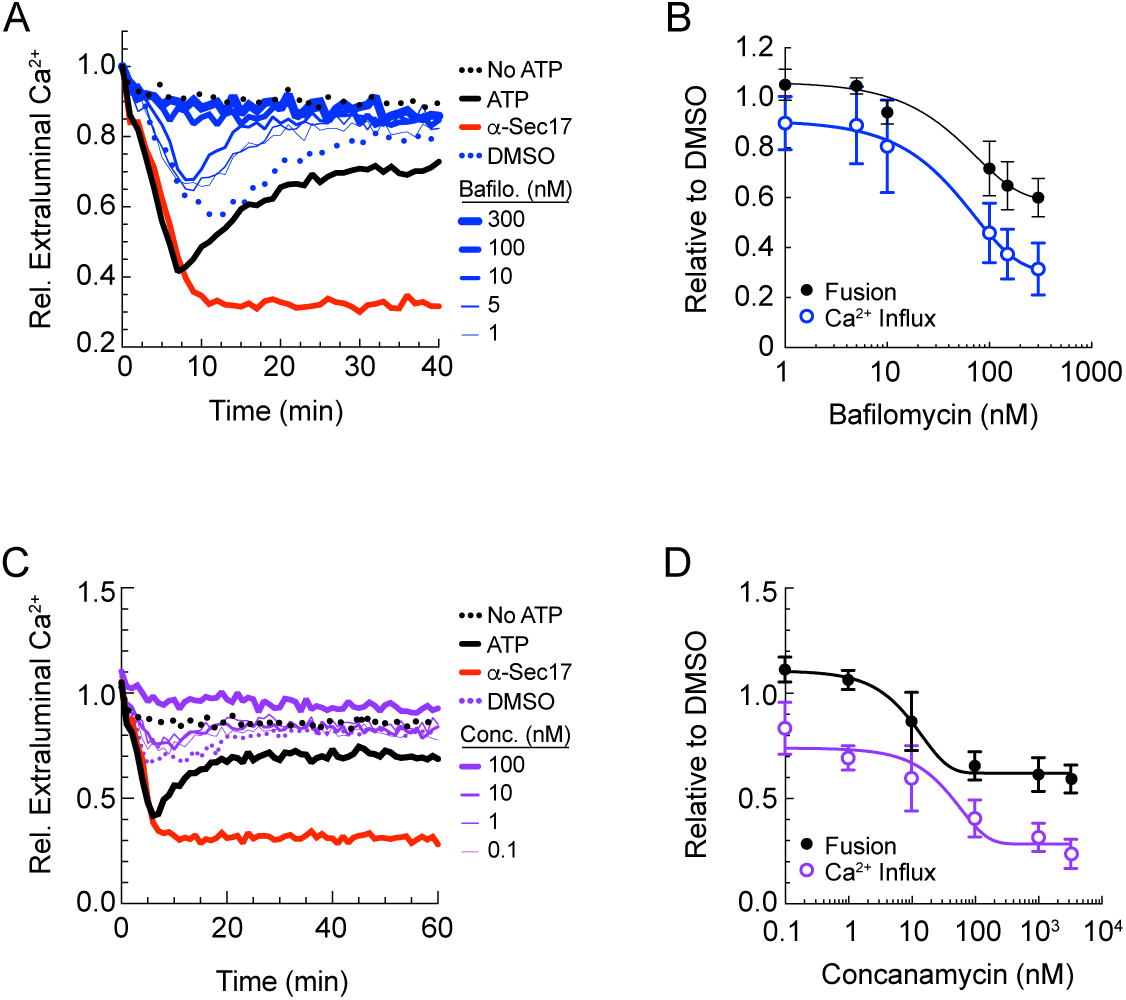
Inhibition of V-ATPase function blocks Ca^2+^ transport. **(A)** Wild type and vacuoles were used in Ca^2+^ transport assays. Reactions were incubated with a dosage curve of Bafilomycin A1 (Bafilo.) or carrier (DMSO). Control reactions were in the presence or absence of ATP or ATP and anti-Sec17 IgG to block SNARE priming. Reactions were incubated for 40 min and Cal520 fluorescence readings were taken every 30 seconds. Fluorescence was normalized to the start of the assay set to 1. **(B)** Average of multiple experiments shown in panel A. In parallel, vacuole fusion reactions were performed in the presence of a dose curve of Bafilomycin A1 **(C)** Ca^2+^ transport assays were performed as in panel A. Instead of Bafilomycin, reactions were incubated with a dose curve of Concanamycin A (Conc.). **(D)** Average of multiple experiments shown in panel C. In parallel, vacuole fusion reactions were performed in the presence of a dose curve of Concanamycin A. Error bars are S.E.M. (n=3).

## Discussion

In this study we present data showing that altered PI(3,5)P_2_ levels proportionately affects V-ATPase activity. Elevating PI(3,5)P_2_ through the hyperactive *fab1*^*T2250A*^ mutation increased V-ATPase activity, whereas blocking Fab1 activity with Apilimod or through kinase dead *fab1*^*EEE*^ mutation decreased V-ATPase function (Banerjee et al., 2019). PI(3,5)P_2_ levels not only modulate V-ATPase function, but as our previous studies showed, changes in PI(3,5)P_2_ affect Ca^2+^ transport through activation of the Ca^2+^ ATPase Pmc1 (Miner et al., 2020). Taken together, these data led to the hypothesis that V-ATPase activity and Ca^2+^ transport are reciprocally regulated through PI(3,5)P_2_ production.

The interdependence of H^+^ and Ca^2+^ transport was shown by testing the effect of changing the concentration of one element on the transport of the second. First, we showed that adding CaCl_2_ to H^+^ pumping assays blocked AO quenching, while reducing free Ca^2+^ with EGTA markedly enhanced AO quenching activity. In comparison, inhibiting the V-ATPase with Bafilomycin A1 or Concanamycin A each inhibited Ca^2+^ uptake. These data support the notion that PI(3,5)P_2_ can affect both H^+^ and Ca^2+^ transport across the vacuole membrane.

Although the connection between PI(3,5)P_2_ and the transport of these ions is strong, it is not the only factor that can inversely control these mechanisms. Ca^2+^ flux can be influenced by other pathways, most notably through the activity of phospholipase C (PLC) on PI(4,5)P_2_, which produces inositol trisphosphate (IP_3_) and diacylglycerol (DAG). During vacuole fusion Plc1 acts on PI(4,5)P_2_ and blocking this through gene deletion or with chemical inhibitors leads to an arrest in vacuole fusion (Jun et al., 2004, Mayer et al., 2000). The inhibitory effect of blocking Plc1 function is attributed to the decrease in the fusogenic lipid DAG, inhibition of Ca^2+^ efflux, as well as altering actin distribution on the vacuole surface (Jun et al., 2004, Karunakaran et al., 2012). The role of DAG in vacuole fusion is further shown by the effects of deleting the DAG kinase Dkg1. Vacuoles lacking Dgk1 not only accumulate DAG but have augmented fusion capacity (Miner et al., 2017). Others have shown that members of the TRPC family of Ca^2+^ channels can be directly affected by DAG or its analog OAG (1-oleoyl-2-acetyl-sn-glycerol) (Venkatachalam et al., 2003). They showed that increased DAG prevents TRPC activation through protein kinase C (PKC) signaling and that TRPC activity was restored by PKC inhibitors. Combined with the effect of PI(3,5)P_2_ on these functions, it is evident that the local lipid environment plays a crucial role in regulating ion transport.

With the interdependence of H^+^ and Ca^2+^ transport on yeast vacuoles, the question arises of whether the channels themselves physically interact? Based on several reports looking at mammalian counterparts, the answer is likely yes. In murine cells, L-type Ca^2+^ channels have been shown to physically interact with the G2 subunit of the H^+^-ATPase (Gao and Hosey, 2000). Others have reported that the R-type Cav2.3 Ca^2+^ channel interacts with the G1-subunit of the V-ATPase and found that Bafilomycin A1 reduces Ca^2+^ transport (Radhakrishnan et al., 2011). In yeast, the addition of an antibody against the Vph1 component of the V_O_ complex blocks Ca^2+^ efflux and vacuole homotypic fusion (Bayer et al., 2003). While not shown directly, it is possible that the antibody against Vph1 physically interfered with the interaction between the V-ATPase and the Ca^2+^ channel. We also found that the Ca^2+^ pump Pmc1 physically interacts with a protein complex that includes Vph1 and the SNARE Nyv1 (Miner et al., 2020, Takita et al., 2001). Based on these studies, we can hypothesize that the physical interaction of transporters is likely a major component of their reciprocal regulation.

Another facet of this paradigm is a direct role of Ca^2+^ or H^+^ ions on each other’s transporters. For the effect of Ca^2+^ on the V-ATPase, we propose that the cation could exert its effects by physically altering membrane fluidity, which in turn negatively affects H^+^ pumping. Other roles for Ca^2+^ release could be interdependently linked with V-ATPase function. Excess extracellular Ca^2+^ blocks acid secretion and H^+^ current in murine osteoclasts by inhibiting V-ATPase at the plasma membrane (Sakai et al., 2006). This idea is supported by multiple studies showing that Ca^2+^ indeed reduces membrane fluidity. This occurs through a direct mechanism by binding to anionic sites on the lipid bilayer, or indirectly by stimulating the expression of membrane enzymes that alter fatty acid composition to vary acyl chain saturation, and in turn affect lipid packing and membrane fluidity. For example, Ca^2+^ reduces the membrane fluidity of brush border and basolateral plasma membrane of intestinal cells (Brasitus and Dudeja, 1986). In rat adipocytes, investigators have shown that increased cytoplasmic Ca^2+^ decreases the fluidity of the plasma membrane inner leaflet (Sauerheber et al., 1980). This effect was reversed by EGTA, indicating that the effect was direct and not due to signaling. In megakaryocytes, increased Ca^2+^ in the cytoplasm released as part of signal transduction decreases lateral mobility (i.e. fluidity) of the membrane (Schootemeijer et al., 1994). The relationship between Ca^2+^ and membrane fluidity can also be observed using artificial membranes. Shah and Schulman showed that Ca^2+^ interacted with phosphatidic acid (PA) preferentially to phosphatidylcholine (PC). Ca^2+^ directly interacts with the free phosphate on PA (Shah and Schulman, 1967). In comparison, PC forms internal salt bridge between its choline and phosphate to preclude a direct interaction with Ca^2+^ even though the dipole made by the PC headgroup moves Ca^2+^ towards the phosphate. Ca^2+^ can also bind phosphatidylserine to form rigid aggregates to alter local membrane fluidity (Ohnishi and Ito, 1973).

How does the V-ATPase affect Ca^2+^ transport? While a direct mechanistic link has not been established for this, there are many examples that are consistent with the notion that the V-ATPase can regulate Ca^2+^ homeostasis as part of membrane trafficking. In *Drosophila*, loss of V-ATPase subunits inhibits the completion of autophagic flux (i.e. final degradation of cargo by pH activated enzymes) yet allows the fusion of autophagosomes with lysosomes (Mauvezin et al., 2015). Bafilomycin inhibits both autophagosome-lysosome fusion as well as pH dependent cargo degradation. The former was thought to be due to the direct inhibition of SERCA by Bafilomycin. Autophagosome-lysosome fusion is stimulated by the activation SERCA and its depletion phenocopies the effects of Bafilomycin on autophagosome-lysosome fusion. Together this in agreement with the idea that the V-ATPase and Ca^2+^ pumps work together.

The fact that the V-ATPase performs functions in addition to H^+^ translocation has been shown by many studies. For instance, in *Drosophila* neurons, the a1 subunit V100 (yeast Vph1) of the V_O_ complex interacts with Ca^2+^-loaded Calmodulin that is needed for eye development (Zhang et al., 2008). In mammalian cells the V-ATPase recruits the small GTPase Arf6 and its nucleotide exchange factor ARNO from the cytosol to endosomal membranes through interaction with the c-ring and a-2 subunits, respectively (Hurtado-Lorenzo et al., 2006). These interactions play a role in endolysosomal degradative pathway. The c-ring in neurons binds the SNARE Synaptobrevin to reduce neurotransmitter release through SNARE-dependent fusion of synaptic vesicles and the plasma membrane (Di Giovanni et al., 2010). In osteoclasts, the d2 subunit promotes osteoclast fusion independent of pH changes caused by V-ATPase (Lee et al., 2006). Finally, the R-type Ca^2+^ channel Cav2.3 binds with G1 subunit of V-ATPase to regulate Ca^2+^ currents (Radhakrishnan et al., 2011). Many more examples like these exist to illustrate that the V-ATPase can physically interact with other proteins to affect a variety of pathways.

In conclusion, this study shows that V-ATPase activity and Ca^2+^ flux can inversely affect each other’s function. While the mechanism by which this occurs remains to be elucidated we can add that their interdependence on the vacuole is coupled with the production of PI(3,5)P_2_ under isotonic conditions. These connections begin to unveil a more complicated network of interactions that integrate the composition of the membrane with ion homeostasis.

## Materials and Methods

### Reagents

Soluble reagents were dissolved in PIPES-Sorbitol (PS) buffer (20 mM PIPES-KOH, pH 6.8, 200 mM sorbitol) with 125 mM KCl unless indicated otherwise. Anti-Sec17 IgG (Mayer et al., 1996), and Pbi2 (Slusarewicz et al., 1997) were prepared as described previously. DiC8-PA (1,2-dioctanoyl-*sn*-glycero-3-phosphate) diC8-PI3P (1,2-dioctanoyl-phosphatidylinositol 3-phosphate), and diC8-PI(3,5)P_2_ (1,2-dioctanoyl-phosphatidylinositol 3,5-bisphosphate) were purchased from Echelon Inc. Apilimod, Verapamil, and Bafilomycin A1 were from Cayman Chemical and dissolved in DMSO. Acridine orange and FCCP (Carbonyl cyanide-4-(trifluoromethoxy) phenylhydrazone were purchased from Sigma and dissolved in DMSO.

### Vacuole Isolation and In-vitro fusion assay

Vacuoles were isolated as described (Haas et al., 1994). *In vitro* fusion reactions (30 µl) contained 3 µg each of vacuoles from BJ3505 and DKY6281 backgrounds, reaction buffer 20 mM PIPES-KOH pH 6.8, 200 mM sorbitol, 125 mM KCl, 5 mM MgCl_2_), ATP regenerating system (1 mM ATP, 0.1 mg/ml creatine kinase, 29 mM creatine phosphate), 10 µM CoA, and 283 nM Pbi2 (Protease B inhibitor). Fusion reactions were incubated at 27°C for 90 min and Pho8 activity was measured in 250 mM Tris-HCl pH 8.5, 0.4% Triton X-100, 10 mM MgCl_2_, and 1 mM *p*-nitrophenyl phosphate. Pho8 activity was inhibited after 5 min by addition of 1 M glycine pH 11 and fusion units were measured by determining the *p*-nitrophenolate produced by detecting absorbance at 400 nm.

### Proton Transport Assay

The proton pumping activity of isolated vacuoles was performed as described by others with some modifications (Müller et al., 2003). *In vitro* H^+^ transport reactions (60 µl) contained 20 µg vacuoles from BJ3505 backgrounds, fusion reaction buffer, 10 µM CoA, 283 nM Pbi2, and 15 µM of the H^+^ probe acridine orange. Reaction mixtures were loaded into a black, half-volume 96-well flat-bottom plate with nonbinding surface. ATP regenerating system or buffer was added, and reactions were incubated at 27°C while acridine orange fluorescence was monitored. Samples were analyzed in a fluorescence plate reader (POLARstar Omega, BMG Labtech) with the excitation filter at 485 nm and emission filter at 520 nm. Reactions were initiated with the addition of ATP regenerating system following the initial measurement. After fluorescence quenching plateaus we added 30 µM FCCP to collapse the proton gradient and restore acridine orange fluorescence. This shows that changes in fluorescence were not due to denaturing of acridine orange by any of the tested reagents or regimens.

### Calcium Efflux

Vacuole lumen Ca^2+^ was measured as described (Miner et al., 2016, Miner and Fratti, 2019, Sasser et al., 2012). *In vitro* Ca^2+^ transport reactions (60 µl) contained 20 µg vacuoles from BJ3505 backgrounds, fusion reaction buffer, 10 µM CoA, 283 nM Pbi2, and 150 nM of the Ca^2+^ probe Cal-520 dextran conjugate MW, 10,000 (AAT Bioquest). Reaction mixtures were loaded into a black, half-volume 96-well flat-bottom plate with nonbinding surface. ATP regenerating system or buffer was added, and reactions were incubated at 27°C while Cal-520 fluorescence was monitored. Samples were analyzed using a fluorescence plate reader with the excitation filter at 485 nm and emission filter at 520 nm. Reactions were initiated with the addition of ATP regenerating system following the initial measurement. The effects of inhibitors on efflux were determined by the addition of buffer or inhibitors immediately following Ca^2+^ influx. Calibration was done using buffered Ca^2+^ standards (Invitrogen).

### Fluorescence microscopy

In vivo vacuole staining with quinacrine and FM4-64 was carried out with log phase yeast grown in YPD broth to an A600 of ∼0.6. Cells were washed with HEPES-KOH, pH 7.6 and stained with 200 µM quinacrine and 5 µM FM4-64 for 10 min at 30°C in the same buffer. Cells were then washed, suspended in YPD and incubated for 1 h at 30°C to chase the FM4-64 to the vacuole. Cells where then washed, suspended in HEPES-KOH, pH 7.6 with 2% glucose, mounted on slides and examined by fluorescence microscopy. Images were acquired using a Zeiss Axio Observer Z1 inverted microscope equipped with an X-Cite 120XL light source, Plan Apochromat 63X oil objective (NA 1.4), and an AxioCam CCD camera. Quinacrine was visualized using a 38 HE EGFP shift-free filter set and FM4-64 was visualized with a 42 HE CY 3 shift-free filter set.

### Statistical analysis

All statistical analysis was calculated using unpaired t-tests. P values of ≤ 0.05 were considered significant.

## Acknowledgments

This research was supported by a grant from the National Institutes of Health (R01-GM101132) and the National Science Foundation (MCB 1818310) to RAF.

## Author contributions

GEM, DAR-K, CZ and RAF, Conception and design, Data analysis and interpretation, Drafting or revising the article; GEM, DAR-K, CZ and KDS Acquisition of data, data analysis.

## Data availability

All data generated or analyzed during this study are included in this published article.

## Abbreviations

AO: acridine orange
DAG: diacylglycerol
FCCP: Carbonyl cyanide-4-(trifluoromethoxy) phenylhydrazone
PA: phosphatidic acid
PI: phosphatidylinositol
PI3P: phosphatidylinositol 3-phosphate
PI(3,5)P_2_: phosphatidylinositol 3,5-bisphosphate
PKC: protein kinase C
PLC: phospholipase C
SNARE: soluble *N-*ethylmaleimide-sensitive factor attachment protein receptor
TRP: transient receptor potential
YPD: yeast extract, peptone, dextrose

